# Balanced Permeability Index: a multi-parameter index for improved in-vitro permeability

**DOI:** 10.1101/2023.12.01.569478

**Authors:** Dahlia R. Weiss, Javier L. Baylon, Ethan D. Evans, Anthony Paiva, Gerry Everlof, Jingfang Cutrone, Fabio Broccatelli

## Abstract

The optimization of passive permeability is a key objective for orally available small molecule drug candidates. For drugs targeting the central nervous system (CNS), minimizing P-gp mediated efflux is an additional important target for optimization. The physicochemical properties most strongly associated with high passive permeability and lower P-gp efflux are size, polarity and lipophilicity. In this study, a new metric called the Balanced Permeability Index (BPI) was developed that combines these three properties. The BPI was found to be more effective than any single property in classifying molecules based on their permeability and efflux across a diverse range of chemicals and assays. The BPI can also be used to guide optimization in non-traditional small molecule modalities, such as protein degraders, which often lie outside of traditional small molecule space. BPI is easy to understand, allowing researchers to make decisions about which properties to prioritize during the drug development process.

---

Human effective permeability across the jejunum membrane has been quantitatively linked to the extent of absorption in human oral experiments for highly soluble drugs.^1^ This relationship has enabled the use of certain cell lines, such as Cancer coli (Caco-2) cells, as suitable tools to rank order marketed drugs based on their jejunum permeability.^2, 3^ Madin-Darby canine kidney (MDCK) cell lines have also been increasingly used in the pharmacokinetics (PK) field over the last two decades to assess passive permeability in a high-throughput assay.^4, 5^ Similar to Caco-2 cells, measurements of passive permeability in MDCK cells are well correlated with human jejunum permeability and bioavailability.^6^ Additionally, MDCK cells can be transfected with P-glycoprotein (P-gp/MDR1) and/or breast cancer resistance protein (BCRP) to assess the risk of efflux mediated by specific membrane transporters that impairs central nervous system (CNS) disposition.^7^

The availability of well validated and robust in vitro assays provides drug discovery scientists with tools to build rational structure activity relationship with respect to oral and CNS absorption. Importantly, the resulting in vitro permeability datasets have allowed linking passive permeability and P-gp mediated efflux to a small number of physicochemical properties, such as molecular weight (MW), heavy atom count (HAC), polarity, number of hydrogen bond donors/acceptors, lipophilicity (e.g., LogD), charge and rigidity.^8-12^ These physicochemical properties have been used as molecular descriptors in machine learning (ML) models have shown promising accuracy at predicting permeability and efflux. However, these models are not readily interpretable, making it difficult to understand how a given result was reached and complicating the assessment of the applicability domain of the predictions. ^13^ To recover interpretability while maintaining predictive power, one strategy is to combine physicochemical properties and well-validated cutoffs into simpler rules. This approach was popularized by Pfizer’s CNS multiparameter optimization model (CNS-MPO) and remains a powerful and intuitive tool for guiding chemistry to design permeable molecules.^14, 15^

It is important to recognize that the areas of chemical space that are more likely to result in optimal absorption may or may not be available to explore based on the therapeutic target and modality. For example, Chen et al. analyzed the distribution of size, polarity, and lipophilicity across different permeability bins (high, medium, and low) using the Genentech compound collection. The analysis demonstrated that physicochemical properties associated with poor permeability (such as size, high polarity, and low lipophilicity) were often correlated with P-gp recognition.^12^ Donovan et al reported over 20K compounds in the Bayer collection for which permeability measurements were available through the Caco-2 assay.^16^ The analysis of those compounds demonstrated an increased tolerated calculated LogD (cLogD) range as the MW is lowered, with a preferred cLogD of 3-3.5 for all MW ranges.

In this study, we present a new Balanced Permeability Index (BPI) that combines three metrics most often associated with permeability: size, polarity, and lipophilicity. Using a large and diverse compound collection in the Bristol-Myers Squibb (BMS) Company, we show that this novel, simple, and interpretable index classifies molecules based on their permeability and efflux class better than any one of its individual components alone. The guidelines resulting from the interpretation of BPI are in line with previously published work: molecules are made smaller, less polar, and more lipophilic, leading to better permeability. An additional learning from BPI is that high polarity (typically linked to low permeability and high efflux) can be balanced by the addition of lipophilic moieties as long as molecule size is kept in check. This can inform non-traditional small molecule projects (e.g., protein degraders) for which the starting point for compounds optimization may lie outside of the boundaries traditionally employed in MPO scores and medicinal chemistry rules of thumb.

We define the experimental Balanced Permeability Index (BPI) as:

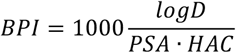

Where logD is the partition coefficient characterizing lipophilicity, PSA is polar surface area and HAC is heavy atom count. To investigate the effectiveness of BPI in predicting compounds with high in vitro permeability, we first turned to the dataset published by Chen et al.^12^ Although the chemical structures were not available as part of the released dataset, the authors reported topological polar surface area (TPSA), HAC and experimentally determined logD at pH 7.4, and measured permeability in two Genentech engineered MDCK cell lines. We were therefore able to validate the ability of the BPI to enrich high permeability compounds, defined as gMDCKI A-to-B Papp > 10 × 10^−6^ cm/s and compounds that are not P-gp efflux substrates, defined as gMDCKI-MDR1 Efflux Ratio < 3. Our initial analysis compared the ability of BPI, TPSA and logD to distinguish those compounds using receiver operating characteristic (ROC) plots (Figure 1A-1B). In this dataset, BPI is better able to enrich the low efflux compounds when P-gp mediated efflux is considered. On the other hand, the added utility of BPI in enriching passive permeability when compared to TPSA is limited in this dataset.

**Figure 1.**
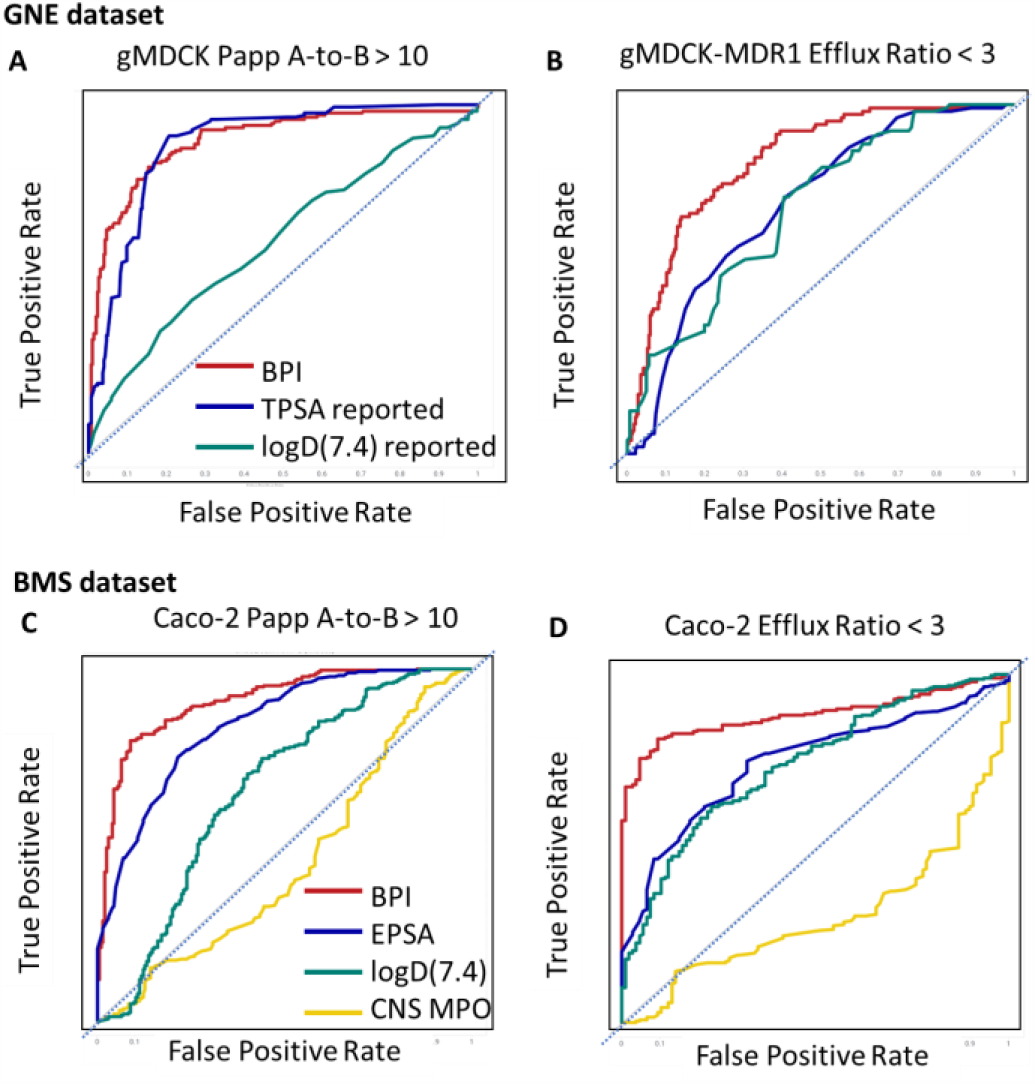
Comparison of the ability of BPI against other parameters to distinguish permeable compounds. ROC plots for: (A-B) Enrichment of permeable Genentech (GNE) compounds gMDCK Papp A-to-B > 10 × 10^−6^ cm/sec^2^ and gMDCK-MDR1 ER < 3 reported in ^12^ using the calculated TPSA (green), LogD_7.4_ (blue) and BPI (red) calculated from those values. (C-D) Enrichment of highly permeable BMS compounds Caco-2 Papp A-to-B > 10 × 10^−6^ cm/sec^2^ and Caco-2 ER < 3 using the measured EPSA (green), LogD_7.4_ (blue) and BPI (red) calculated from those, and compared to the Pfizer CNS MPO (yellow) described in ^15^

Next, we analyzed the BMS chemical compound collection. As in previous reports, we did not rely on calculated logP or logD, instead measuring lipophilicity with a high-throughput chromatographic LogP assay at pH 7.4 (LogD_7.4_). The HPLC LogP assay was similar to previously reported utilizing a retention time-based approach where compounds, along with a set of calibration standards with known LogP values, were analyzed with an isocratic HPLC-UV/MS method with a mobile phase of 60% methanol and 40% ammonium acetate (5 mM) on a YMC C18 ODS-A column (2.1 × 50 mm, 5 μm) at a flow rate of 0.5 mL/min. The compounds’ LogP values were then calculated based on comparison of their retention times to those of the standards (p-methoxyphenol, p-cresol, 1-naphthol, thymol, diphenyl ether and hexachlorobenzene).^17^

TPSA is calculated solely based on the 2-dimensional structure of the molecule, which may not accurately represent the true 3-dimensional polar surface area of a molecule in solution. In our analysis, we utilized the experimentally determined exposed polar surface area (EPSA). The original EPSA assay was described by Goetz et al, and the data in this work was measured by using our inhouse EPSA assay that was modified for higher throughput.^18^ For simple compounds TPSA is a reliable indicator of compound polarity in solution and in those cases may be a useful tool for assessing passive permeability, as demonstrated in the Genentech data set. However, for drug-like molecules that have the potential for intra-molecular hydrogen bonds, TPSA is not an accurate reflection of the EPSA measured in solution. In fact, a study of more than 11,450 EPSA data points in the BMS compound collection found no correlation to the calculated TPSA value, with an R^2^ value of 0.07 (data not shown).

We evaluated a total of 2393 compounds with available data for both EPSA and LogD_7.4_. To ensure the reliability of our analysis, we applied strict cutoffs to the permeability assay data, limiting the recovery between 80% and 120%. This approach minimized the risk of poor recovery affecting the interpretation of the results. Our analysis included 690 compounds with measured passive permeability data in a MDCK cell line with knocked-out endogenous canine MDR1 P-gp efflux transporter (MDCK-KO), and 481 molecules with permeability data in the Caco-2 cell line.^19^ We excluded compounds for which efflux could not be accurately quantified due to low permeability outside of the quantifiable range. Highly permeable molecules were defined as those with Caco-2 Papp A-to-B permeability > 10 × 10^−6^ cm/sec^2^, Caco-2 Efflux Ratio < 3 and MDCK-KO Papp A-to-B permeability > 10 × 10^−6^ cm/sec^2^, and low permeability is defined as Caco-2 Papp A-to-B permeability < 1.5 × 10^−6^ cm/sec (the lowest level of detection), Caco-2 Efflux Ratio > 10 and MDCK KO Papp A-to-B permeability < 10^−6^ cm/sec^2^.

The analysis of BMS compounds revealed that the use of the BPI parameter provides an advantage over relying solely on EPSA and LogD_7.4_ alone to classify compounds in the Caco-2 high permeability and low efflux bin (Figure 1C-1D). While each parameter alone can identify permeable compounds, BPI leads to a better enrichment and higher early enrichment. We also compared our results to the Pfizer CNS MPO, which did not enrich the permeable compounds in the Caco-2 cell line in this collection. It is important to note that most compounds in the collection were not specifically designed to be brain penetrant. To evaluate the predictive ability of each classifier, we used the area under the ROC curve, a common metric used to characterize the predictive ability of a given classifier. Our results demonstrated that BPI is more predictive than EPSA and LogD_7.4_ alone across all the permeability and efflux assays (Table 1).

**Table 1.**
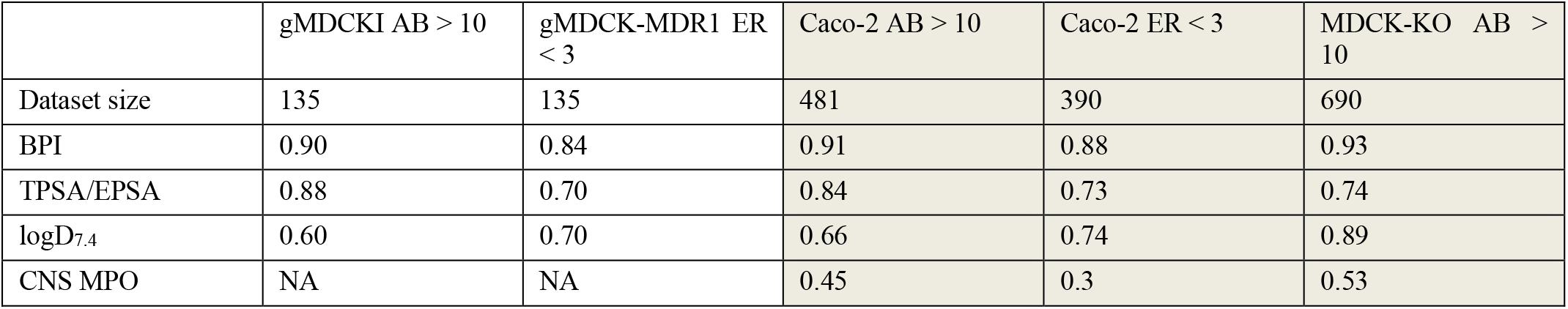
ROC-AUC for enrichment of highly permeable and low efflux compounds. The gMDCKI and gMDCK-MDR1 datasets in the white columns are reported in ^12^, while the highlighted Caco-2 and MDCK-KO columns are proprietary datasets.

The distributions of BPI, EPSA, logD_7.4_ and HAC for the high, medium, and low permeability/efflux bins in the BMS collection are shown in Figure 2. The distributions were found to be statistically different (Kruskal-Wallis one-way ANOVA test, p<0.05) in most cases. One exception is the MDCK-KO Papp A-to-B permeability, where the difference with the LogD_7.4_ distribution was not statistically significant (p>0.05). These findings further support the idea that BPI is more predictive of passive permeability and low efflux than relying solely on the physicochemical properties included in the index. It is important to note that we did not filter the molecules based on size, pKa, or chemical class, and our analysis included molecules with different sizes, polarities, and lipophilicities, with ranges of HAC between 15-65, EPSA between 40-320, and logD between -1 to 6. The lack of compound-related exclusion criteria, in addition to the translatability across chemical spaces, cell lines and datasets from different companies, strongly suggests that the BPI metric can generalize beyond the data used in the analysis.

**Figure 2.**
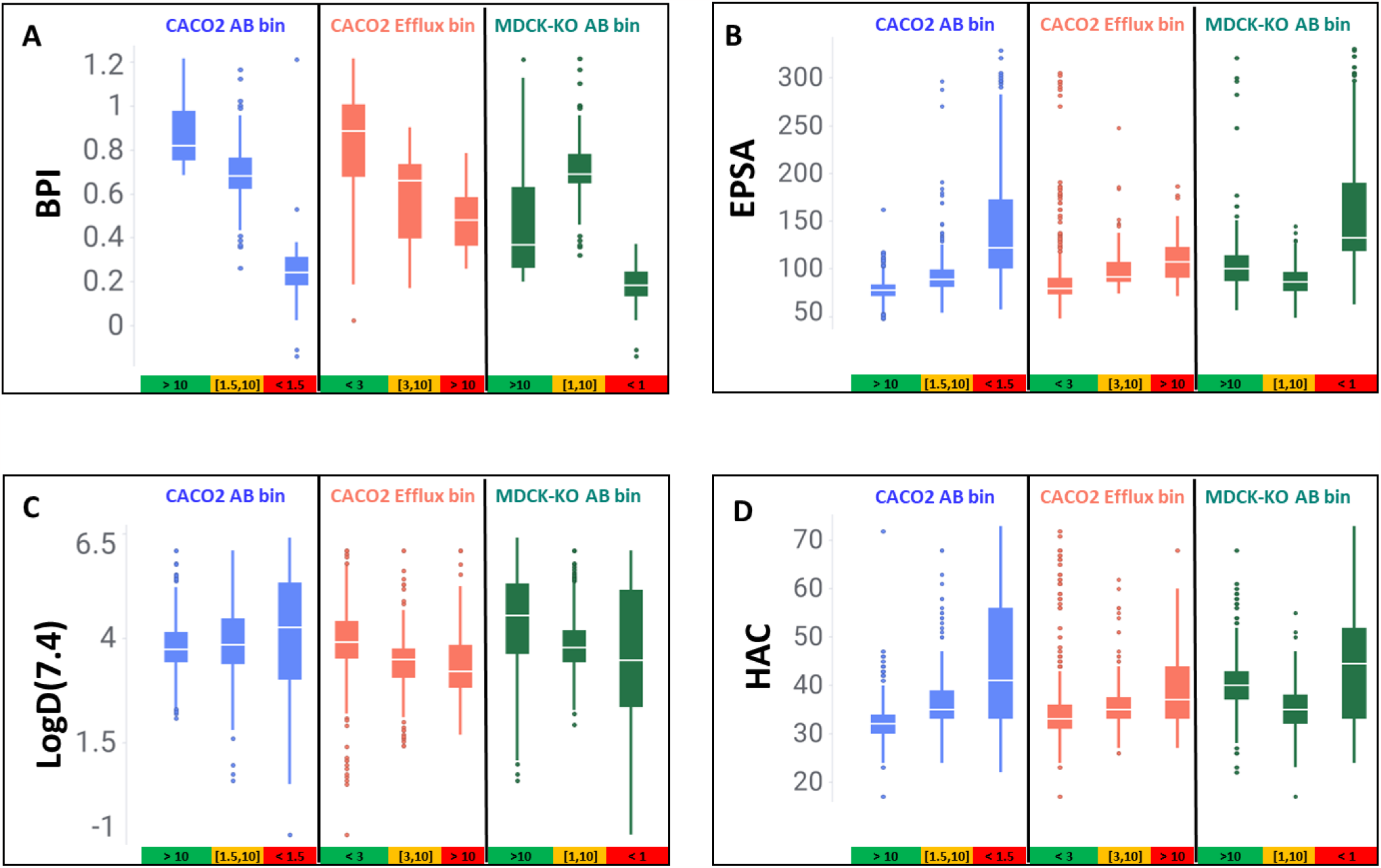
Distribution of BPI and other physicochemical properties across different permeability and efflux ranges. The distribution of (A) BPI, (B) logD, (C) EPSA and (D) HAC split by permeability bins as measured by the Caco-2 A-to-B forward permeability (blue boxes), the Caco-2 efflux ratio (pink boxes) and the MDCK-KO A-to-B forward permeability (yellow boxes). Compounds are binned according to permeability as follows: highly permeable molecules are defined as Caco-2 Papp A-to-B permeability > 10 × 10^−6^ cm/sec^2^, Caco-2 Efflux Ratio < 3 and MDCK-KO Papp A-to-B permeability > 10 × 10^−6^ cm/sec^2^, and low permeability is defined as Caco-2 Papp A-to-B permeability < 1.5 × 10^−6^ cm/sec^2^ (the lowest level of detection), Caco-2 Efflux Ratio > 10 and MDCK KO AB permeability < 10^−6^ cm/sec^2^, and the middle bin falling between those two ranges.

It is suggested that cut-offs be associated with BPI to provide tangible objectives during compound optimization. The recommended BPI cutoff may vary based on the experimental assays and therapeutic indication. However, based on the available data, tentative guidelines have been provided (as shown in Figure 3). The BPI bins are colored by measured Caco-2 Papp A-to-B permeability and efflux ratio, which supports a guidance of BPI >1 for programs looking for orally dosed small molecule compounds, and BPI>1.5 for programs looking for exposure in the central nervous system, which requires a higher permeability and lower efflux.

**Figure 3.**
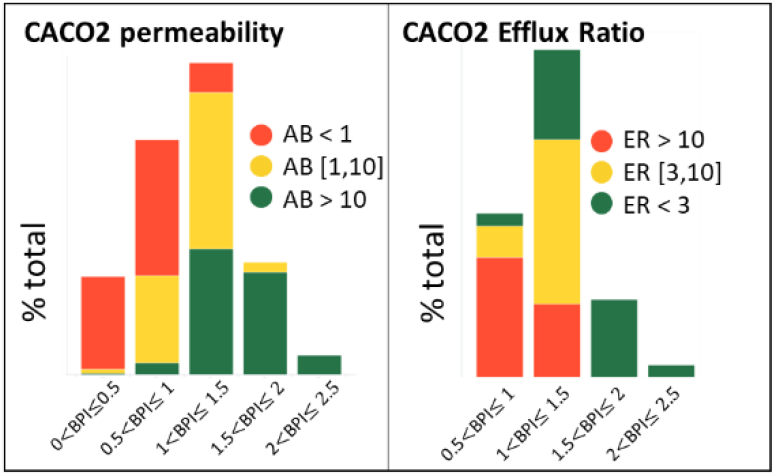
Caco-2 permeability binned by BPI in the BMS collection dataset. Binning of BPI provides some guidelines to enrich compounds with high permeability and low efflux (green boxes in both panels): BPI >1 for orally dosed small molecules, and BPI>1.5 for molecules targeting central nervous system exposure, which requires high permeability and low efflux.

To further validate the guidelines for BPI, a correlation between BPI and free brain-to-plasma partition from rodent experiments was investigated.^20^ The brain and plasma protein binding were either derived from equilibrium dialysis or Transil binding assays selected by PK experts based on the scaffold specific characteristics. In the analysis, free brain to plasma ratio (Kpu,u) and calculated BPI (based on measured EPSA and internal machine learning models for LogD_7.4_) showed a high degree of correlation with R^2^=0.7 (Figure 4a). For this specific analysis, the choice of using calculated LogD_7.4_ was a function of the data availability, not a preference towards calculated parameters. Notably, the Kpu,u dataset includes compounds from 3 different therapeutic projects and six distinct chemotypes. The correlation between Kpu,u and calculated BPI further suggests that this new metric can generalize across different chemotypes, as previously mentioned in the analysis of datasets from different companies. In this analysis, the calculated BPI differentiates between brain penetration classes (low, moderate, higher based on Kpu,u cut-offs of 0.1 and 0.3) with minimal overlap, as shown in Figure 4b.

**Figure 4.**
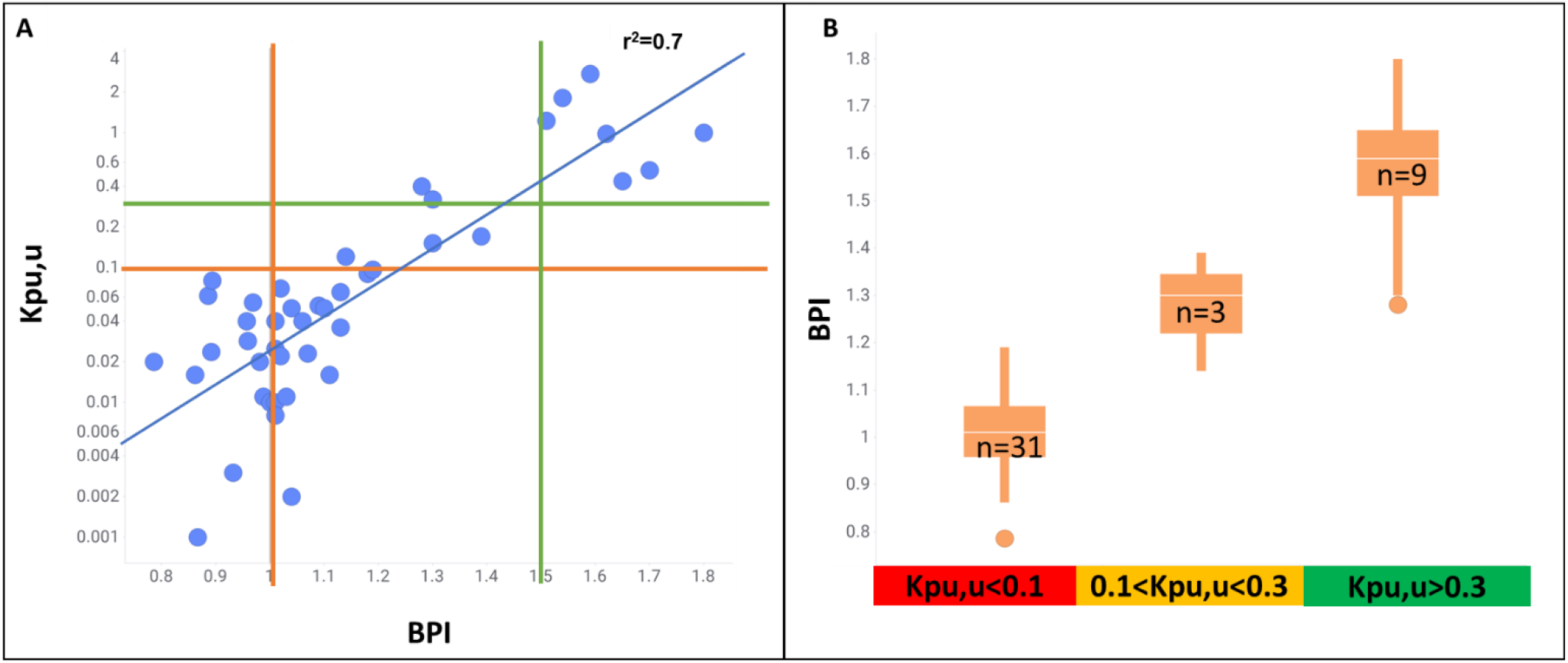
BPI is highly indicative of compound ability to partition into the CNS. (A) The log(Kpu,u) is highly correlated to BPI (R^2^=0.7, N=44). Compounds originated from three programs and six chemotypes across the BMS portfolio, all with an aim to achieve CNS penetrance. (B) The BPI distribution across low, medium and high Kpu,u bins can be used to classify compounds. In this case, a BPI>1 is recommended to increase the likelihood of finding medium and high permeability compounds, while BPI>1.5 ensures highly permeable compounds.

In conclusion, the BPI is a new metric that combines size, polarity, and lipophilicity. It is better able to distinguish highly permeable, low efflux compounds than any of the constituent properties alone in both a published dataset and across the proprietary compound collection. The BPI is easily understood and can guide compound design towards smaller, less polar, and more lipophilic compounds. This versatile metric can be applied to a wide range of chemical space, including novel small molecule modalities such as protein degradation, which lie outside the typical physicochemical property space. Adoption of BPI as a multiparameter score can guide chemistry into better in-vitro permeability space, raising the likelihood that compounds will show desirable oral and CNS availability in-vivo.

## ACKNOWLEDGMENT

We would like to acknowledge the work of the Lead Discovery and Optimization ADME team for permeability measurements, and the Pharmaceutical Candidate Optimization team.

## ABBREVIATIONS

BPI: Balanced Permeability Index
CNS: central nervous system
P-gp: P-glycoprotein
Caco-2: Cancer coli
PK: pharmacokinetics
MDCK: Madin-Darby canine kidney
MDCK-KO: MDCK knock out
BCRP: breast cancer resistance protein
MW: molecular weight
ML: machine learning
MPO: multiparameter optimization
BMS: Bristol-Myers Squibb Company
HAC: heavy atom count
TPSA: Topological polar surface area
EPSA: exposed polar surface area
ROC: receiver operating characteristic
AUC: area under curve
ER: efflux ratio
Kpu,u: unbound brain-to-plasma drug partition coefficient

## Notes

### Competing Interest Statement

The authors have declared no competing interest.

